# An indocyanine green-based liquid biopsy test for circulating tumor cells for pediatric liver cancer

**DOI:** 10.1101/2023.07.03.547557

**Authors:** Andres F. Espinoza, Pavan Kureti, Roma H. Patel, Saiabhiroop R. Govindu, Bryan W. Armbruster, Martin Urbicain, Kalyani R. Patel, Dolores Lopez-Terrada, Sanjeev A. Vasudevan, Sarah E. Woodfield

## Abstract

**Background and Aims:** Hepatoblastoma (HB) and hepatocellular carcinoma (HCC) are the most common malignant hepatocellular tumors seen in children. The aim of this work was to develop a liquid biopsy test for circulating tumor cells (CTCs) for these tumors that would be less invasive and provide information about the real-time state of tumors in response to therapies.

**Methods:** For this test, we utilized indocyanine green (ICG), a far-red fluorescent dye that is used clinically to identify malignant liver cells in the body during surgery. We assessed ICG accumulation in cell lines with fluorescence microscopy and flow cytometry. For our CTC test, we developed a panel of liver tumor-specific markers, ICG, Glypican-3 (GPC3), and DAPI and tested this panel with cell lines and non-cancer control blood samples. We then used this panel to analyze whole blood samples for CTC burden with a cohort of 14 HB and HCC patients and correlated with patient characteristics and outcomes.

**Results:** We showed that ICG accumulation is specific to liver cancer cells, compared to non-malignant liver cells, non-liver solid tumor cells, and non-malignant cells and can be used to identify liver tumor cells in a mixed population of cells. Experiments with the ICG/GPC3/DAPI panel showed that it specifically tagged malignant liver cells. With patient samples, we found that CTC burden from sequential blood samples from the same patients mirrored the patients’ responses to therapy.

**Conclusions:** Our novel ICG-based liquid biopsy test for CTCs can be used to specifically count CTCs in the blood of pediatric liver cancer patients.

**Impact and implications:** This manuscript represents the first report of circulating tumor cells in the blood of pediatric liver cancer patients. The novel and innovative assay for CTCs shown in this paper will facilitate future work examining the relationship between CTC numbers and patient outcomes, forming the foundation for incorporation of liquid biopsy into routine clinical care for these patients.

**Graphical abstract:** Overview of novel liquid biopsy test for circulating tumor cells for pediatric liver cancer. Figure made with Biorender.

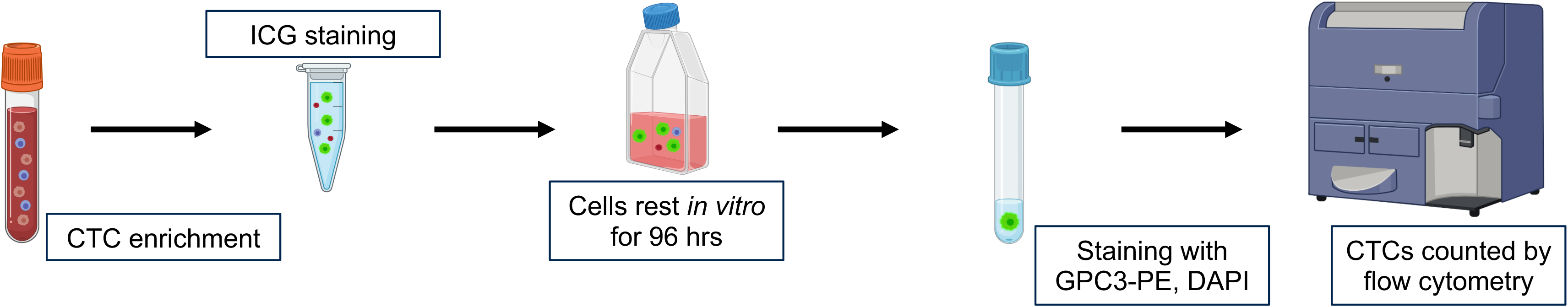

## INTRODUCTION

Hepatoblastoma (HB) is the most common malignant hepatocellular tumor seen in children and generally affects individuals younger than 5 years of age with a less than 50% overall survival (OS) in patients that present with multifocal, metastatic, or treatment refractory disease^1–3^. Notably, HB has the fastest rising incidence of all pediatric solid tumors^4^ due to its association with premature birth, maternal environmental exposures, and cancer predisposition syndromes^5,6^. Current frontline therapy for HB includes aggressive surgery, either resection or orthotopic liver transplant, with cisplatin/doxorubicin-based chemotherapy. Despite receiving intense treatment, children with high-risk HB attain less than 50% 5-year event free survival and suffer many permanent toxicities related to therapy^1,7–10^. Pediatric hepatocellular carcinoma (HCC) has a dismal 5-year OS rate of approximately 28%^11^, although patients with resectable disease achieve a much better OS rate of 70-80%^12^. Medical therapy for pediatric HCC is mainly limited to a chemotherapy regimen based on cisplatin and doxorubicin (PLADO), although kinase inhibitor sorafenib has been tested as well^13^. The most meaningful impact on HCC outcomes has been in the field of interventional radiology where chemo- or radio-embolization can be used to prolong survival^14^. Fibrolamellar carcinoma (FLC) is a very rare subtype of HCC that occurs in adolescents and young adults that has a 5-year overall survival rate of approximately 50% for children^15,16^. Surgical resection is the only curative treatment for this disease, and there are no standard of care systemic therapies for FLC^15^.

Generally, pediatric patients with liver tumors are initially diagnosed by imaging and assessment of serum levels of Alpha-fetoprotein (AFP), followed by biopsy for tissue diagnosis and subtype classification. Treatment plans are individualized based on patient age, histopathological and imaging information, and blood AFP levels^2^. During therapy, longitudinal clinical monitoring of patients is completed with imaging and blood AFP levels. Patients are deemed disease-free when tumors are no longer visible by current imaging modalities and AFP has dropped to normal levels.

Unfortunately, current strategies of matching individual cases to effective therapeutic strategies are suboptimal. HB and pediatric HCC are very heterogeneous tumors, and, with limited tumor material from biopsies, not all tumor sub-clones may be captured. In addition, AFP levels can be misleading as some tumors do not secrete AFP. In addition, infants, particularly premature babies, may have high levels of AFP because of ongoing development of the liver^17^. For children, repeated imaging can increase future oncologic risk and cause neurological damage as they are exposed to radiation and repeat sedation. Standard imaging modalities do not have the resolution needed for detection of individual tumor cells, and patients may harbor one or a few tumor cells at the end of therapy^18^. Clinical tests with the ability to detect “minimal residual disease” in the form of circulating tumor cells (CTCs) may aid in identifying such patients before they progress to visible or detectable recurrence or metastases.

Liquid biopsy assays for CTCs circumvent many of these challenges. Liquid biopsy involves a peripheral whole blood draw, followed by assessment of the blood for tumor material, including CTCs and circulating tumor DNA (ctDNA). Sampling the blood for tumor cells gives a readout of the most aggressive clones present that are already in the process of disseminating^19^. In addition, sequential sampling of CTCs can provide information about the evolution of tumor cells as they change in response to therapies^19^. These cells provide a precise evaluation about the real-time state of the tumor at diagnosis or as the patient undergoes therapy^19^. The major challenge of developing liquid biopsy assays for CTCs is how to unambiguously identify these cells in the blood; these tumor cells are very rare, approximately 1 in 10^6^ leukocytes^20^. Both marker-based and physical property-based protocols to isolate CTCs have been validated. The key to marker-based protocols is using markers that are common to all tumor cells, even if these cells evolve as they enter the blood and disseminate.

To address these challenges in our development of a liquid biopsy test for CTCs, we harnessed the power of indocyanine green (ICG), a far-red fluorescent dye that is established to accumulate in tumor cells with a liver origin. ICG is currently validated for use clinically during surgery to identify liver tumor cells throughout the body, including cells that represent intrahepatic primary liver tumors or extrahepatic disease, including lung metastasis^21–27^. To validate the ICG^+^ cells as liver tumor cells, we chose a second marker that is specific to HB and HCC tumor cells, Glypican-3 (GPC3). GPC3 was shown to be elevated in a study of 60 HB tumor samples and negative in adjacent benign liver parenchyma, illustrating the specificity of this marker for HB^28^. In addition, GPC3 is established to be overexpressed in HCC^29^. This specificity of GPC3 for hepatocellular cancer cells has led to the development of GPC3-targeting chimeric antigen receptor (CAR) T cell trials that are currently being evaluated in clinical trials enrolling pediatric and adult patients with liver tumors.

In this paper, we describe a novel liquid biopsy assay for pediatric liver cancer CTCs. This test is based on the unambiguous identification of cells that express three markers, ICG, Glypican-3 (GPC3), and DAPI with both standard and imaging flow cytometry. Importantly, this test can be used with the most common pediatric primary tumors of the liver, HB and HCC, including the FLC subtype of HCC.

## MATERIALS AND METHODS

### Cells and culture conditions

The HepG2, Huh-6, HepRG, and A549 cell lines were commercially acquired (HepG2 (HB-2065), A549 (CRM-CCL-185): American Type Culture Collection (ATCC), Manassas, VA, USA; Huh-6: Riken Cell Bank, Japan; HepRG: HPRGC10, Invitrogen, Waltham, MA, USA). The SH-SY5Y and 293T cell lines were generously provided by Dr. Jianhua Yang (Baylor College of Medicine, Houston, TX, USA). All widely available cell lines were grown in Eagle’s Minimum Essential Medium (EMEM, Lonza, Allendale, NJ, USA) supplemented with 10% heat-inactivated fetal bovine serum (FBS, SAFC Biosciences, Lenexa, KS, USA), 2 mM glutamine (Invitrogen), and 100 units/ml streptomycin/penicillin (Invitrogen).

The HB17 cell line was developed in our laboratory from a patient-derived xenograft (PDX) tumor. Cells from the tumor were grown *in vitro* initially on Matrigel (354230, Corning, Glendale, AZ, USA) in Hepatocyte Culture Medium (CC-3198, Lonza). After 40 passages, the cells were transitioned to standard tissue culture plasticware without Matrigel.

All cells were grown at 37 °C in 5% CO2. All cells were validated with short tandem repeat (STR) DNA profiling yearly. Profiles of widely available cell lines were compared to a database of cell line profiles. The HB17 profile was compared to a profile generated by STR DNA profiling of the paired primary patient sample. All cells were also tested for mycoplasma (MycoAlert, Lonza) every year.

### ICG

ICG was obtained as a powder (United States Pharmacopeia, MD, USA) diluted to 2 mg/ml in water, and then added at 25 μM to cells and patient samples.

### Fluorescent antibodies and other markers

A GPC3 antibody conjugated to the PE fluorophore (clone 1G12 + GPC3/863, NBP2-47763PE, Novus Biologicals, Centennial, CO USA) was used with immunofluorescence and flow cytometry experiments as described at a dilution of 1:100. For living cells, DAPI (cat. no. R37605, NucBlue Live Cell Stain ReadyProbes, Invitrogen) was used according to the manufacturer’s protocol for microscopy experiments. For fixed cells, DAPI (cat. no. R37606, NucBlue Fixed Cell Stain ReadyProbes, Invitrogen) was used for flow cytometry experiments at a dilution of 2 drops per 1 ml.

### Standard flow cytometry

Samples were analyzed with flow cytometry on a standard Symphony instrument (BD Biosciences) using a 637 nm laser to excite ICG and a 780/60 bandpass filter to detect it, a 561 nm laser to excite PE and a 586/15 bandpass filter to detect it, and a 405 nm laser to excite DAPI and a 431/28 bandpass filter to detect it. All cell line samples were gated as follows: (1) forward scatter (FSC) - area versus side scatter (SSC) - area to identify cells (2) FSC - area versus FSC - width to identify single cells; (3) SSC – height versus SSC - width to identify single cells; (4) histogram of DAPI - area to identify DAPI^+^ enucleated cells; and (5) histogram of APC-Cy7 - area to identify ICG^+^/DAPI^+^ cells; (5) histogram of PE – area to identify GPC3-PE^+^/ICG^+^/DAPI^+^ cells (if applicable). All patient samples were gated as follows: (1) FSC - area versus FSC - width to identify single cells; (2) SSC - height versus SSC - width to identify single cells; (3) histogram of DAPI - area to identify ICG^+^/DAPI^+^ enucleated cells; (4) histogram of APC-Cy7 - area to identify ICG^+^ cells; (5) histogram of PE – area to identify GPC3-PE^+^/ICG^+^/DAPI^+^ cells. All samples were run with appropriate positive and negative controls as described. All patient samples were run with the following HepG2 and A549 positive and negative controls: (1) HepG2 negative; (2) HepG2 ICG^+^/GPC3^+^/DAPI^+^; (3) A549 negative; (4) A549 GPC3^+^/DAPI^+^. All data was analyzed with FlowJo version 10.6.2 (Becton Dickinson). For patient samples, collection was performed on the entirety of the sample.

### Imaging flow cytometry

Samples were analyzed with flow cytometry on an Amnis Imagestream X MKII (Luminex) equipped with 405 nm, 488 nm, 561 nm, 633 nm and 785 nm scatter lasers. Collection was performed on the entirety of each sample. Objects were analyzed and gated using IDEAS software 6.3.23.0 as follows: (1) cells were gated using Aspect Ratio and Area parameters for the brightfield channel. First, focused cells were gated using the Gradient_RMS parameter for Channel 1 (Brightfield). Second, single cells were gated using Area versus Aspect Ratio for Channel 1 (Brightfield). (2) DAPI^+^ single cells were gated with a histogram of Intensity of Channel 7 (CCR7/BV421) with the single cell population. (3) DAPI^+^/ICG^+^/GPC3^+^ single cells were gated with a scatter plot of Intensity of Channel 12 (APC-Cy7/APC-ef780/Live Dead NIR) versus Intensity of Channel 3 (PE) with the DAPI^+^ population. Representative images used for figures were exported from the IDEAS image gallery as .tif files and inserted into the manuscript. Raw data files are available upon request. All samples were run with appropriate HepG2 and A549 positive and negative controls: (1) HepG2 negative; (2) HepG2 ICG^+^/GPC3^+^/DAPI^+^; (3) HepG2 ICG^+^; (4) HepG2 GPC3^+^; (5) HepG2 DAPI^+^; (6) A549 negative; (7) A549 GPC3^+^/DAPI^+^.

### Microscopy

Brightfield and fluorescent images of cells were taken on a BZ-X710 All-in-One Fluorescence Microscope (Keyence, Itasca, IL, USA) at the indicated magnifications and scales with ICG and DAPI filters. The objective used was a Nikon 20X PlanFluor objective with a numerical aperture of 0.45. The microscope and acquisition software used was the standard for this microscope. All images were taken at room temperature. Cells were alive in cell culture media and tagged with mCherry, ICG, and DAPI as indicated.

### Patient samples

Samples were collected from patients after informed consent was obtained from their parents or guardians via an Institutional Review Board (IRB)-approved blood collection protocols H-38834 and H-49313. All experiments on patient samples were performed in compliance with the Helsinki Declaration and approved by the Baylor College of Medicine IRB.

### Processing of human whole blood samples

Samples were depleted of cells of an immune origin with the RosetteSep CD45 depletion kit (depletes with antibodies for CD45, CD66, and Glycophorin A, cat. no. 15122, Stem Cell Technologies, Cambridge, MA USA) according to the protocol with SepMate - 15 tubes (cat. no. 85415, StemCell Technologies) and Lymphoprep (cat. no. 1114544, Alere Technologies, San Diego, CA USA). Remaining cells were then washed with ACK lysis buffer (cat. no. 118-156-101, Quality Biological, Gaithersburg, MD USA) to lyse remaining red blood cells. Samples were then incubated with 25 μM ICG for 1 hour at 37°C. After ICG incubation, cells were kept in culture for 96 hours in complete hepatocyte basal medium (cat. no. CC-3198, Lonza). Cells were then fixed in 4% paraformaldehyde (cat. no. 15710, Electron Microscopy Sciences, Hatfield, PA USA) for 30 minutes and stained with GPC3-PE antibody (1:100) for 30 minutes. Cells were incubated with DAPI (see above) prior to flow cytometry experiments.

### GPC3 IHC

Patient primary tumor and lung metastasis samples were fixed in 10% formalin, embedded in paraffin, and mounted on glass slides in the Texas Children’s Hospital clinical pathology laboratory as part of routine clinical care of these patients. Slides were stained with Glypican-3 (1G12, Cell Marque, Rocklin, CA USA). Stained slides were then scanned with a Leica Aperio AT2 slide scanner. The slides were pre-processed to select the bounding area and focus points for each slide. After scanning, the images were saved on a secure hospital server. The GPC3 images shown in the manuscript are representative of the entire samples and were selected from the whole scanned area using QuPath version 0.4.3. GPC3 histology was quantified by a pathologist (K.R.P.) with a standard scoring system of Intensity (I) + Extent (E) = T (Total). Intensity was scored as 0 (absent), 1 (weak), 2 (intermediate), or 3 (strong). Extent was scored as 0 (0), 1 (>0% to <25%), 2 (>25% to <50%), 3 (>50% to <75%), or 4 (>75% to 100%). This information is shown in Table 2, along with histology of each tissue sample (also assessed by a pathologist (K.R.P.)).

### Patient ICG

ICG was utilized during hepatectomy and metastasectomy for pediatric liver tumors as described^21^ at Texas Children’s Hospital. ICG images of primary samples shown in Figures 4, 5, and 6 were obtained from electronic medical records.

### Patient characteristics

We reviewed electronic medical records for the following patient-specific information shown in Table 1: (1) diagnosis, (2) pretreatment extent of disease (PRETEXT) stage^30^, (3) Children’s Oncology Group (COG) stage^7^, (4) risk group^2^, (5) presence of multifocal disease in the liver, (6) presence of vascular invasion (VI), (7) presence of metastasis, and (8) ICG positivity. We reviewed electronic medical records for the following sample-specific information shown in Table 2: (1) time point in care when sample was received, (2) whether chemotherapy had been received by the patient at that time point, (3) AFP level at that time point, and (4) disease burden at that time point. Disease burden was determined by assessing available clinical data to estimate whether tumor burden was increasing or decreasing following therapy. Diagnostic biopsy samples are considered separate from other samples for increasing or decreasing burden. We were blinded to CTC burden when we assessed disease burden. CT images shown in Figures 4, 5, and 6 were taken as part of routine clinical care and were extracted from electronic medical records for this manuscript.

**Table 1.** Table of patient data. N/A for ICG^+^ indicates that ICG was not used during surgery so information about ICG accumulation is not available.

|  | Diagnosis | PRETEXT stage | COG stage | Risk | Multifocal | VI | Metastasis | ICG+ |
| --- | --- | --- | --- | --- | --- | --- | --- | --- |
| HB70/110 | HB | 1 | 1 | VLR | N | N | N | Y |
| HB73 | HB | 4 | 4 | HR | Y | Y | Y | N/A |
| HB77 | HB | 2 | 1 | LR | N | Y | N | Y |
| HB78 | HB | 3 | 1 | IR | Y | N | N | N/A |
| HB80 | HB | 4 | 1 | HR | N | N | Y | Y |
| HB99 | HB | 3 | 4 | HR | N | Y | Y | Y |
| HB102/106 | HB | 3 | 2 | IR | Y | Y | N | Y |
| HB103/113 | HB | 4 | 4 | HR | Y | Y | Y | Y |
| HB108 | HB | 4 | 3 | IR | Y | Y | N | N/A |
| HB109 | HB | 2 | 1 | VLR | N | Y | N | N/A |
| HB112 | HB | 3 | 2 | HR | N | Y | Y | Y |
| HB83/114/116 | HCC | 4 | 4 | HR | Y | Y | Y | Y |
| HB105 | FLC | 3 | 2 | HR | Y | Y | Y | Y |
| HB119 | FLC | 4 | NA | HR | Y | Y | N | N/A |
| HB71 | MH | 2 | NA | NA | N | N | N | N/A |

## RESULTS

### ICG selectively accumulates in liver cancer cells

We first sought to show that ICG selectively accumulates in liver cancer cells *in vitro* by fluorescence microscopy. We optimized dosing of ICG and imaging timing to show positive ICG presence in liver tumor cells with accompanied negative signal in non-tumor and non-liver cells. As shown in Figure 1, after incubation with ICG for 1 hour at a dose of 25 µM, we see clear ICG accumulation in the HepG2, Huh-6, and HB17 HB tumor cells 24 hours later (Figure 1A). At the same dose and time points, we see lack of ICG in the non-tumor HepRG cell line, the non-liver SH-SY5Y and A549 cell lines, and the non-liver and non-tumor 293T cell line (Figure 1A). Results remain consistent at extended intervals of 48, 72, and 96 hours post-incubation with ICG (Figure 1A). Notably, the HepG2 and HB17 cells appear even brighter at the 72 and 96 hour post-incubation intervals. At the same time points, we see low or negative signal in the non-tumor and non-liver cell lines (Figure 1A).

**Figure 1.**
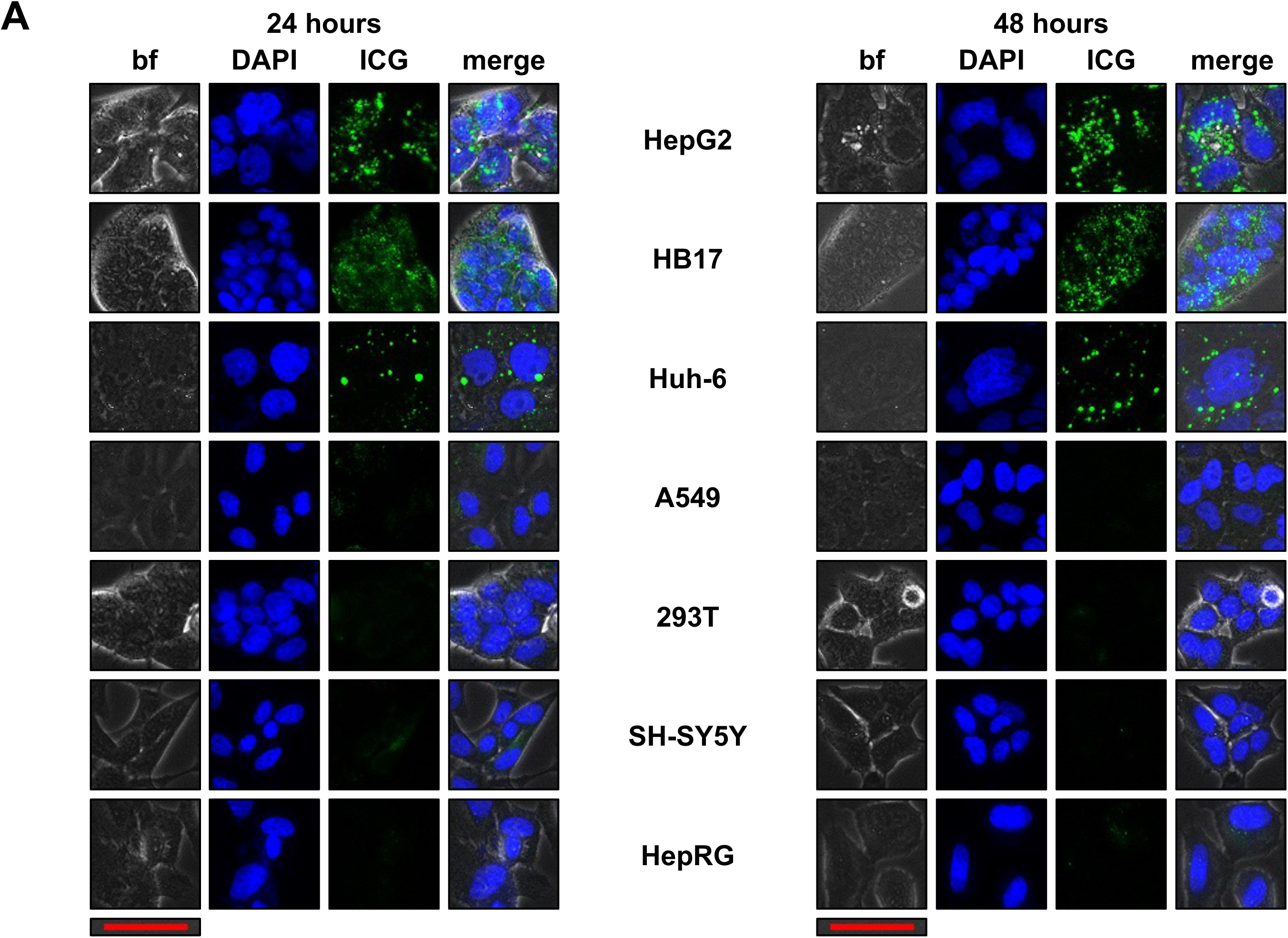
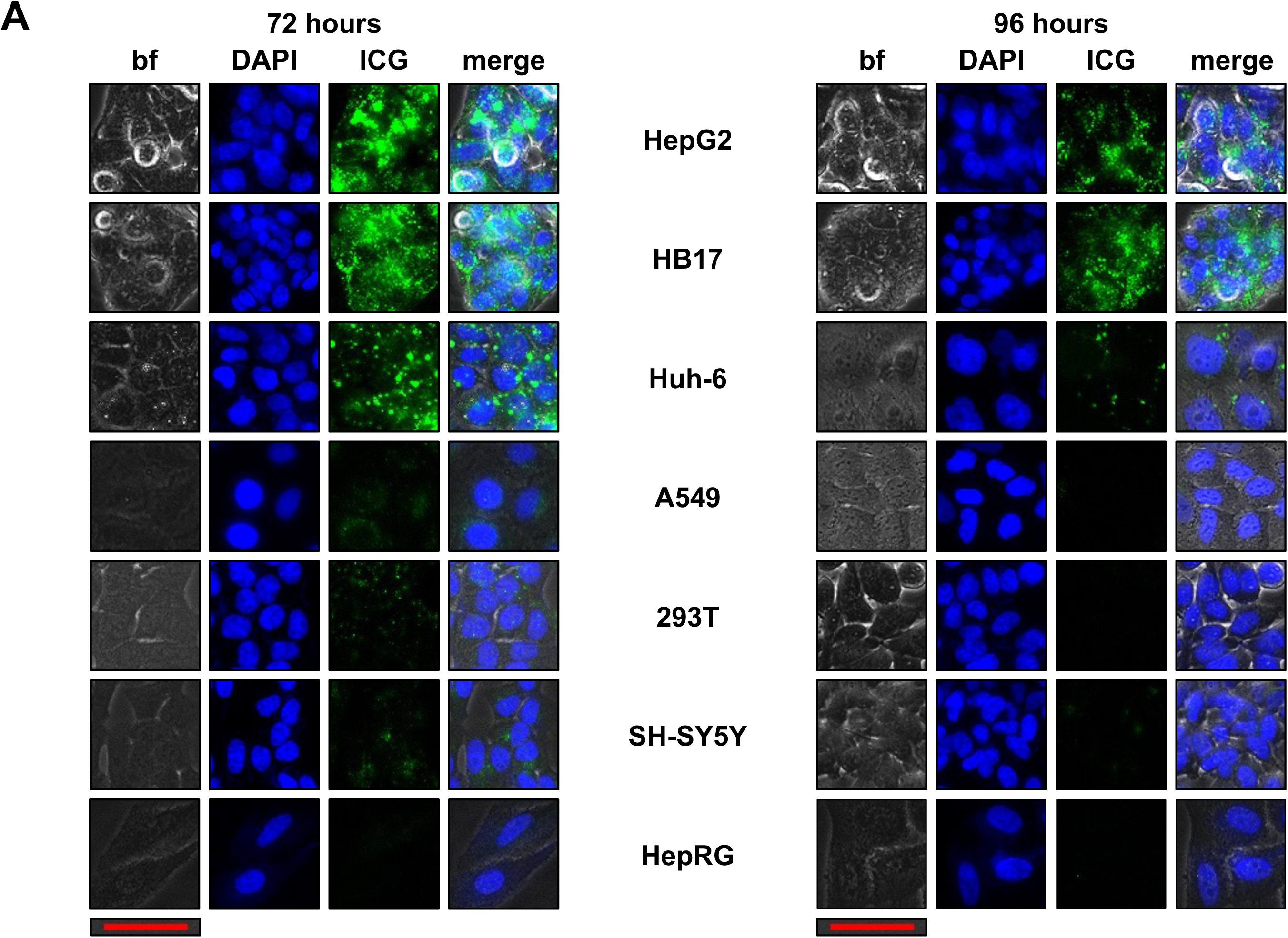
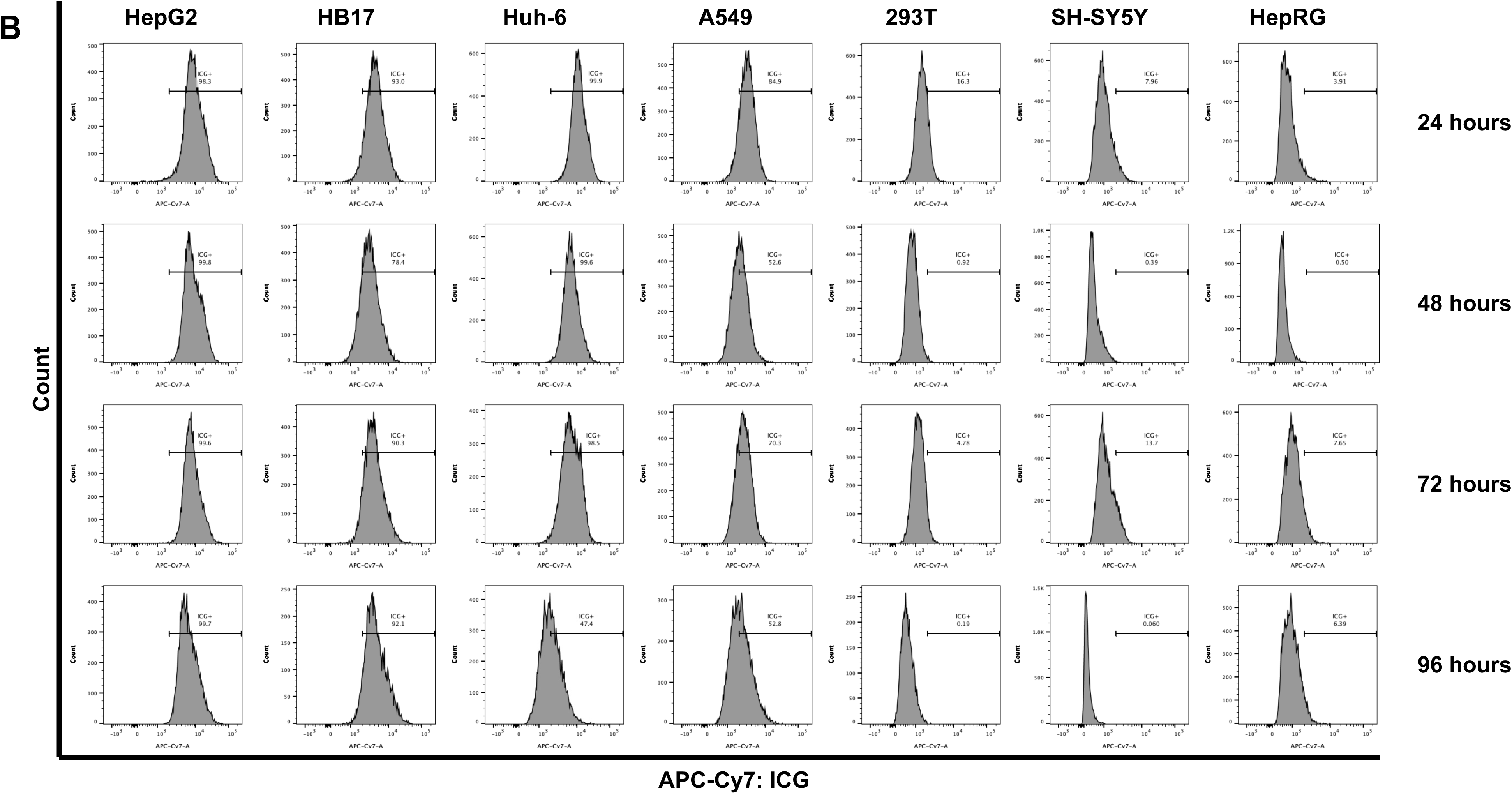
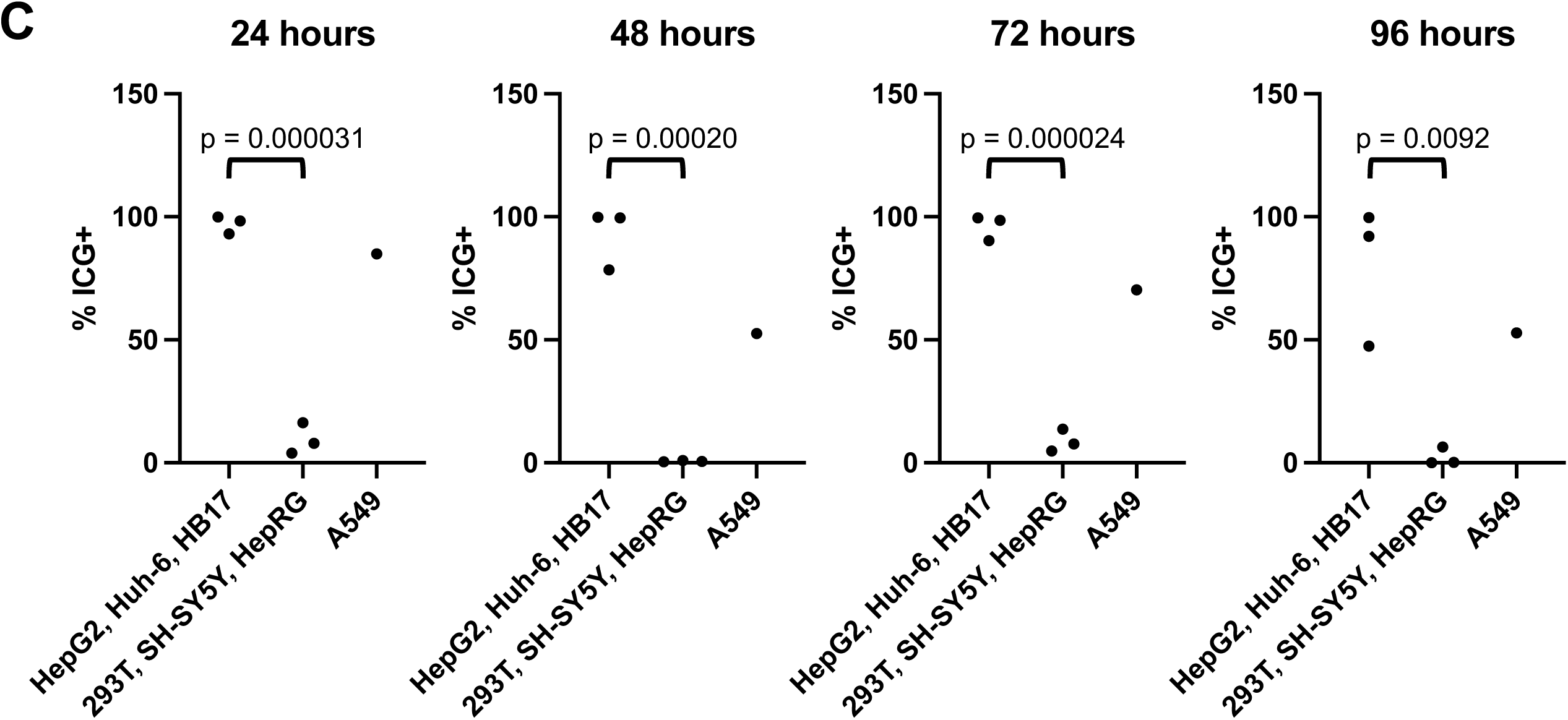
ICG selectively accumulates in liver cancer cells. (A) Cells stained with 25 μM ICG for 1 hour and then imaged 24, 48, 72, and 96 hours after incubation. Cells were also mixed with DAPI to indicate nuclei to show cells present in each image. Scale bar (red) represents 50 μm. (B,C) Cells stained with 25 μM ICG for 1 hour and then analyzed by flow cytometry at 24, 48, 72, and 96 hours after incubation. Histograms of the DAPI^+^/ICG^+^ population in each sample shown in B. Percent positive for ICG^+^ shown in C with statistical significance analyzed by student’s t-test (two tailed). Exposure time 2 seconds for 24 hour time point, 5 seconds for 48, 72, and 96 hour time points.

This ICG positivity can also be analyzed by flow cytometry with APC-Cy7 fluorophore conditions. HepG2, HB17, and Huh-6 cells stained with 25 µM ICG and flowed 24, 48, 72, and 96 hours after incubation show clear high APC-Cy7/ICG signal, while SH-SY5Y, 293T, and HepRG cells stained with the same concentration of ICG and analyzed by flow cytometry at the same time points have a very low signal (Figure 1B,C). By flow cytometry, A549 cells show intermediate ICG accumulation (Figure 1B,C). This data is further analyzed in Figure 1C, showing that malignant liver cells show statistically significant higher levels of ICG fluorescence intensity at all four time points, compared to 293T, SH-SY5Y, and HepRG cells. Taken together, this work shows that ICG accumulation can be used to unambiguously identify tumor cells of a liver origin.

### ICG far-red fluorescent signal can be used to identify liver cancer cells in a mixed population of cells

We further verified that ICG positivity is specific to liver cancer cell lines and not secondary to staining conditions by analyzing mixed populations of cells by fluorescence microscopy. For these experiments, we used HepG2 cells expressing mCherry (HepG2-mCherry). We mixed HepG2-mCherry cells with either 293T or SH-SY5Y cells at a 1:1 ratio and then stained all cells with 25 µM ICG for 1 hour, followed by imaging at 48 and 96 hours after 1 hour incubation with ICG. Figure 2A shows clear positive ICG signal in the mCherry^+^ HepG2 cells while the 293T or SH-SY5Y cells in the same well remain negative.

**Figure 2.**
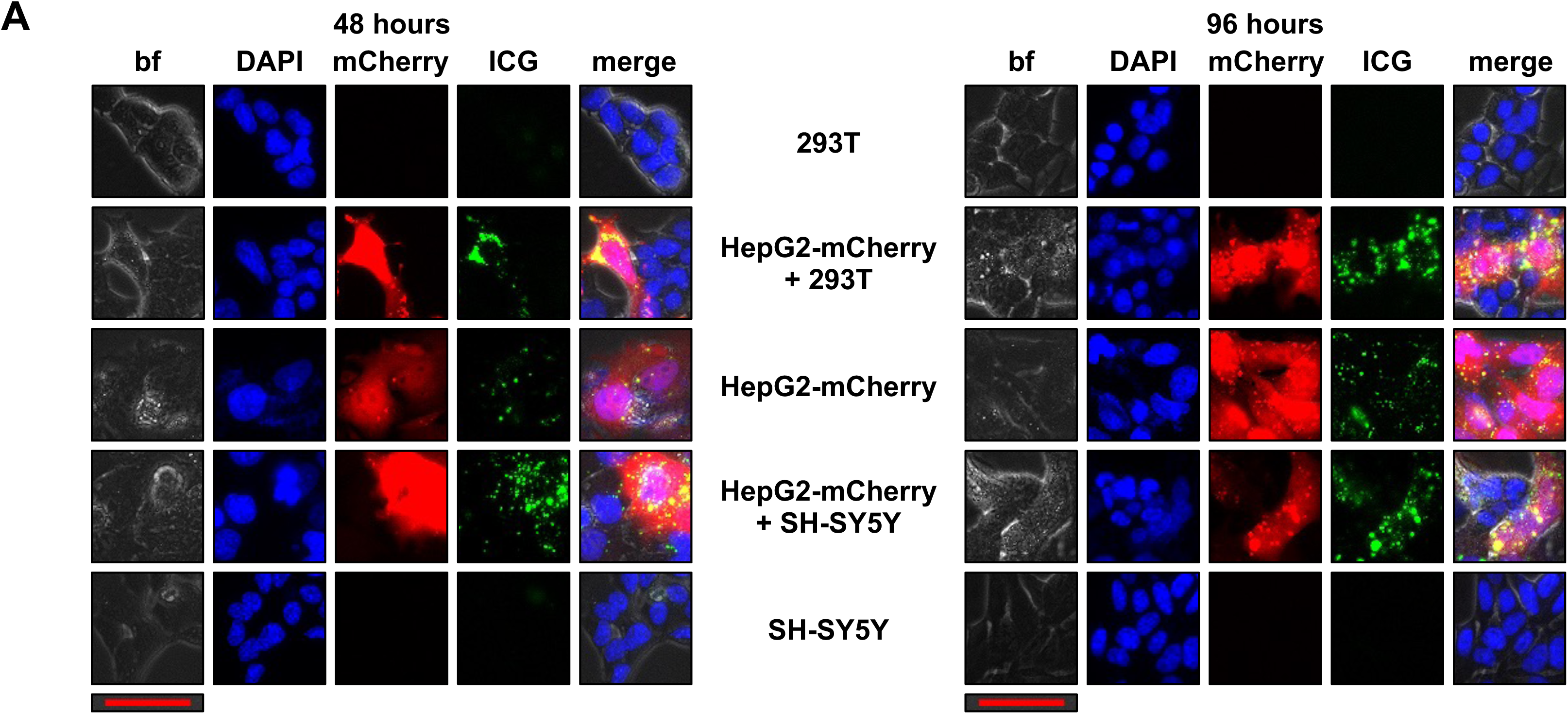
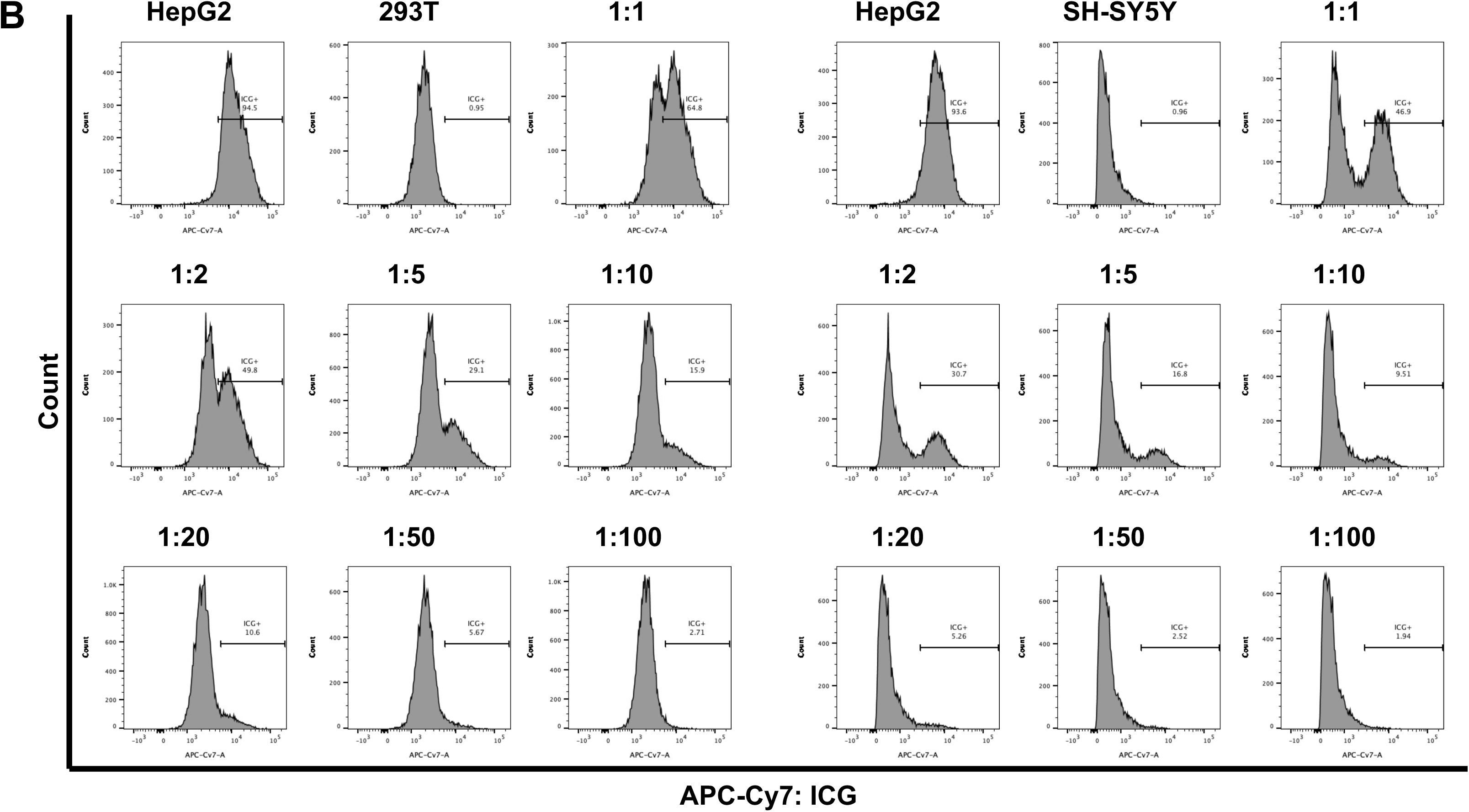
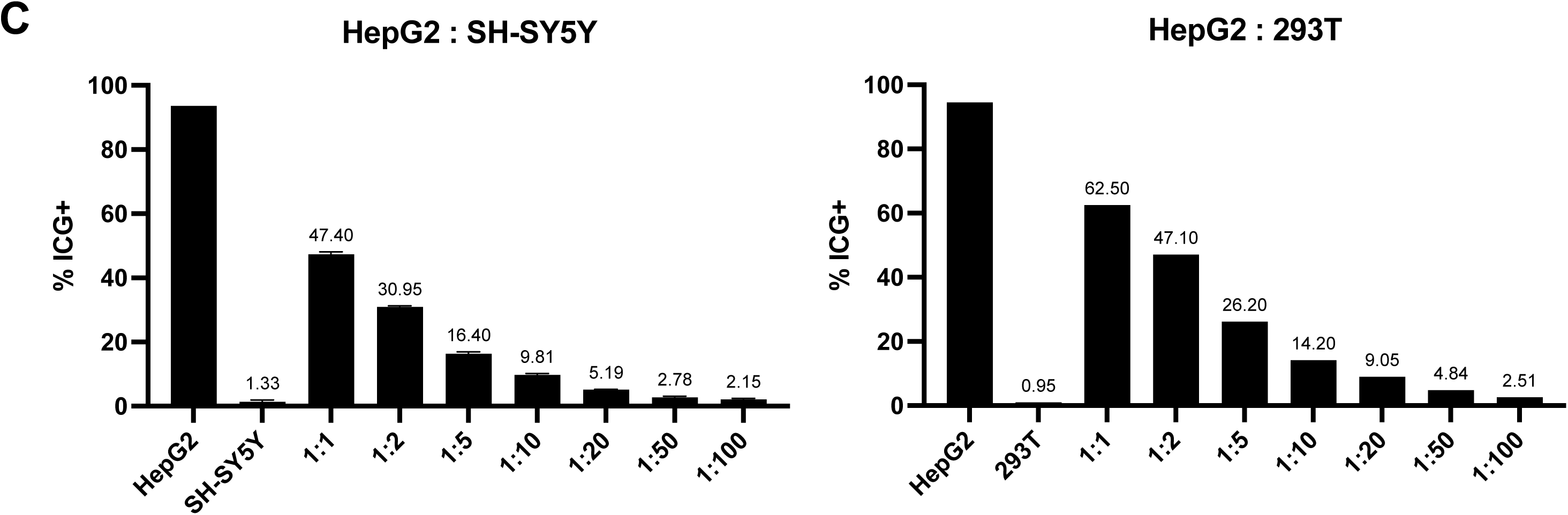
ICG can be used to identify liver cancer cells in a mixed population of cells. (A) HepG2-mCherry cells mixed with 293T or SH-SY5Y cells were stained with ICG for 1 hour then imaged 48 and 96 hours after incubation. Cells were also mixed with DAPI to indicate nuclei to show cells present in each image. Scale bar (red) represents 50 μm. (B,C) HepG2, 293T, and SH-SY5Y cells were stained with ICG for 1 hour and then analyzed by flow cytometry 96 hours after incubation. HepG2 cells were mixed with either 293T or SH-SY5Y cells at the indicated ratios. Histograms of the DAPI^+^/ICG^+^ population in each sample shown in B. Graphs in C show percent ICG^+^ cells in each mixture. Exposure time 5 seconds for both 48 and 96 hour time points.

In addition, this ICG positivity can be used to selectively identify HB tumor cells mixed with 293T or SH-SY5Y cells by flow cytometry. We mixed HepG2 with 293T or SH-SY5Y cells that were both incubated with 25 µM ICG for 1 hour. We mixed them together at a range of ratios as indicated in Figure 2B,C and analyzed them by flow cytometry with APC-Cy7 fluorophore conditions to count ICG positivity at a single 96 hour time point after ICG incubation, the point at which we saw the greatest difference in positive and negative populations. The histograms in Figure 2B show the separation of the two populations of cells at both time points when the cells were mixed at the indicated ratios. The graphs in Figure 2C show that the percent positive for ICG coincides to the ratio of HepG2 cells present in the mix.

### Unambiguous identification of liver tumor cells with a panel of ICG, GPC3, and DAPI

To further validate the liver tumor origin of ICG^+^/DAPI^+^ cells, we added an antibody recognizing human GPC3, which is well established to be specifically expressed in HB^28^. Shown in Figure 3A are histograms showing that 98.6% of DAPI^+^ HepG2 cells are also GPC3^+^. In contrast, all DAPI^+^ A549 cells are GPC3^-^. This data is further analyzed in Figure 3B, showing that liver tumor cells have a statistically significant increase in GPC3 expression compared to A549 cells. Multiple replicates of this experiment are plotted in Figure 3B showing the reproducibility of this experiment.

**Figure 3.**
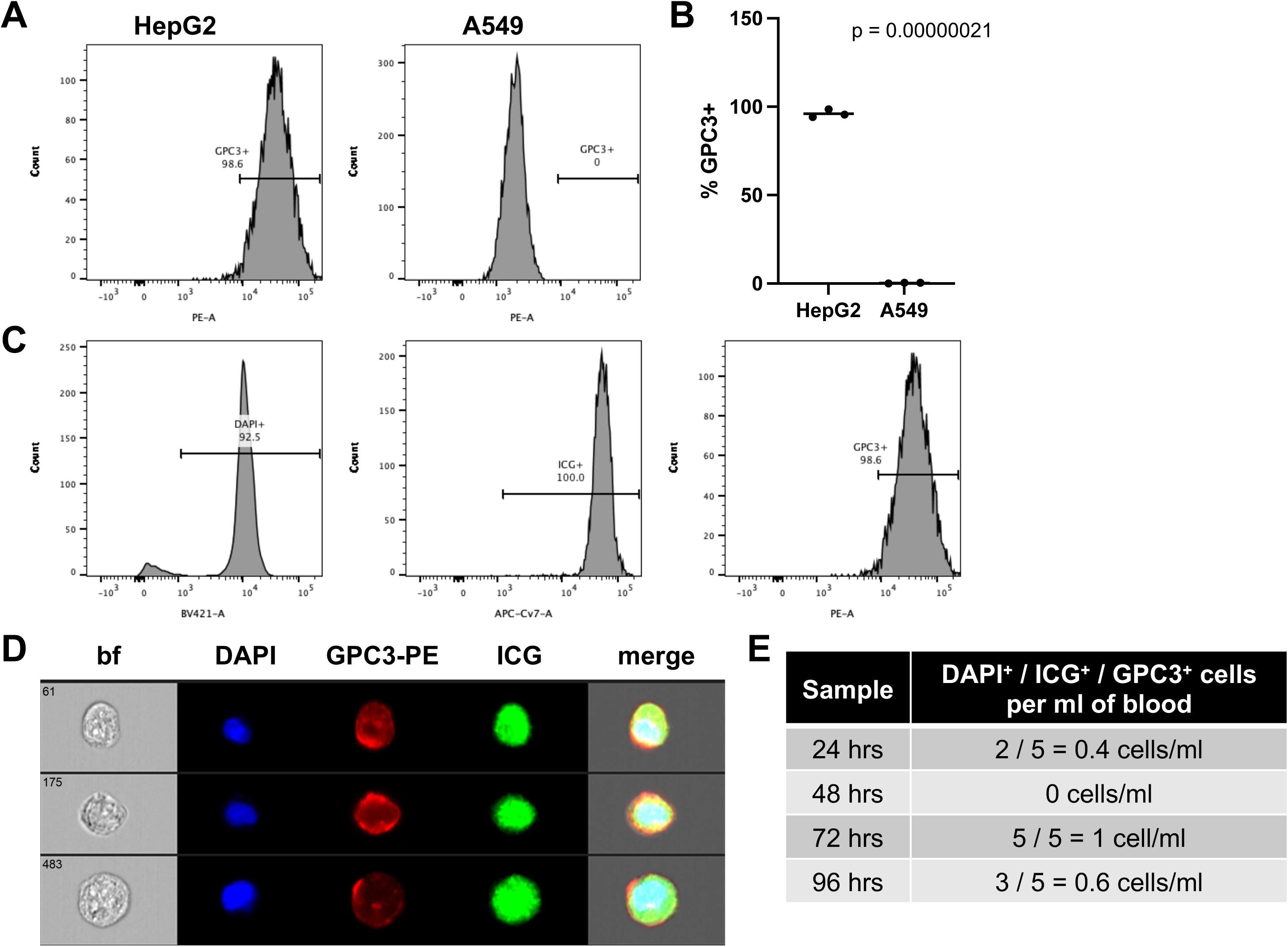
A marker panel of ICG, GPC3, and DAPI can be used to unambiguously identify liver tumor cells. (A,B) Flow cytometry for GPC3 with HepG2 positive control cells and A549 negative control cells shows the specificity of GPC3 for HepG2 cells that express this protein. Histograms of the DAPI^+^/GPC3^+^ population in each sample shown in A. Percent DAPI^+^/GPC3^+^ for each sample graphed in B with statistical significance analyzed by student’s t-test (two tailed). (C) Gating strategy for samples run with the DAPI/ICG/GPC3 panel. Single cells were first separated by gating with FSC and SSC as described in the methods. DAPI^+^/ICG^+^/GPC3^+^ cells were then separated and counted on histograms as follows: (1) DAPI^+^ enucleated cells, (2) ICG^+^ cells, and (3) GPC3^+^ cells. This is shown for a representative HepG2 sample. (D) Images of DAPI^+^/ICG^+^/GPC3^+^ HepG2 cells analyzed by the Amnis ImageStream instrument. (E) DAPI^+^/ICG^+^/GPC3^+^ cells present in non-cancer control blood samples. Blood samples were processed as described in the methods, and samples were stained with ICG for 1 hour, followed by resting in standard cell culture conditions in HBM for the indicated amount of time. Samples were then analyzed by flow cytometry on a BD Symphony instrument for numbers of DAPI^+^/ICG^+^/GPC3^+^ cells. All samples showed 1 cell or fewer per ml of blood.

We developed a panel to unambiguously identify pediatric liver tumor cells that includes ICG, GPC3, and DAPI. After staining, we analyzed the samples by flow cytometry on a BD Biosciences standard flow cytometer and an Amnis ImageStream imaging flow cytometer, both with APC-Cy7, PE, and BV421 lasers. We first tested this panel with the HB cell line HepG2 and the lung cancer cell line A549. Shown in Figure 3C is our gating strategy with our control HepG2 sample analyzed on the Symphony instrument. We used histograms for each channel to independently identify each population as follows: (1) DAPI^+^ cells with BV421-Area, (2) DAPI^+^/ICG^+^ cells with DAPI^+^ cells with APC-Cy7-Area, and (3) DAPI^+^/ICG^+^/GPC3^+^ cells with DAPI^+^/ICG^+^ cells with PE-Area. Importantly, with this strategy, 92.5% of HepG2 cells were DAPI^+^; of those, 100% were ICG^+^; and, of those, 98.6% were GPC3^+^, showing the specificity and sensitivity of the panel for identifying liver cancer cells (Figure 3C). Shown in Figure 3D are our control HepG2 cells flowed on the Amnis ImageStream instrument.

We then tested our panel on non-cancer control whole blood samples. We collected samples of 20 mls total of whole blood to analyze at four time points after collection and incubation with ICG, 24 hours, 48 hours, 72 hours, and 96 hours. First, to decrease the contamination of immune cells in our downstream assays, we depleted samples for immune cells with the RosetteSep CD45 depletion kit (depletes with antibodies for CD45, CD66, and Glycophorin A). We then stained the remaining cells with our panel of ICG, GPC3-PE, and DAPI and analyzed the samples on the Symphony standard flow cytometer. Shown in Figure 3E, no samples showed more than 1 cell/ml of DAPI^+^/ICG^+^/GPC3^+^ cells.

### Unambiguous identification of CTCs with immune cell depletion and a panel of ICG, GPC3-PE, and DAPI

Twenty whole blood samples from a cohort of 15 patients were analyzed with this panel of markers and standard and imaging flow cytometry. These patients represent diagnoses of HB, HCC, and FLC. One patient with a non-malignant tumor, mesenchymal hamartoma (MH) was used as a control. Characteristics for these patients are shown in Table 1. Counts for all of these samples on the standard and imaging flow cytometers, as well as individual sample characteristics, are shown in Table 2. Histological assessment and GPC3 positivity of a sample from each patient is also shown in Table 2. Importantly, correlating these patient and sample characteristics with characteristics, including PRETEXT stage, COG stage, risk group, presence of multifocal disease, presence of vascular invasion, presence of metastasis, chemotherapy at time of sample, AFP at time of sample, and disease burden, did not show any statistically significant correlations (Supplementary Figure 1). We did observe a trend towards an association with higher CTC counts with diagnosis or with increasing disease burdens. For samples that had a CTC count on both the standard and imaging cytometers, we generally saw agreement with an average difference of 12.81 cells/ml and only two samples with a difference greater than 10 cells/ml. As expected, the patient with MH had a very low count of 1.2 cells/ml, similar to the non-cancer control samples. For all patients, we checked GPC3 expression and ICG accumulation to validate the use of our test based on GPC3 and ICG positivity.

**Table 2.**
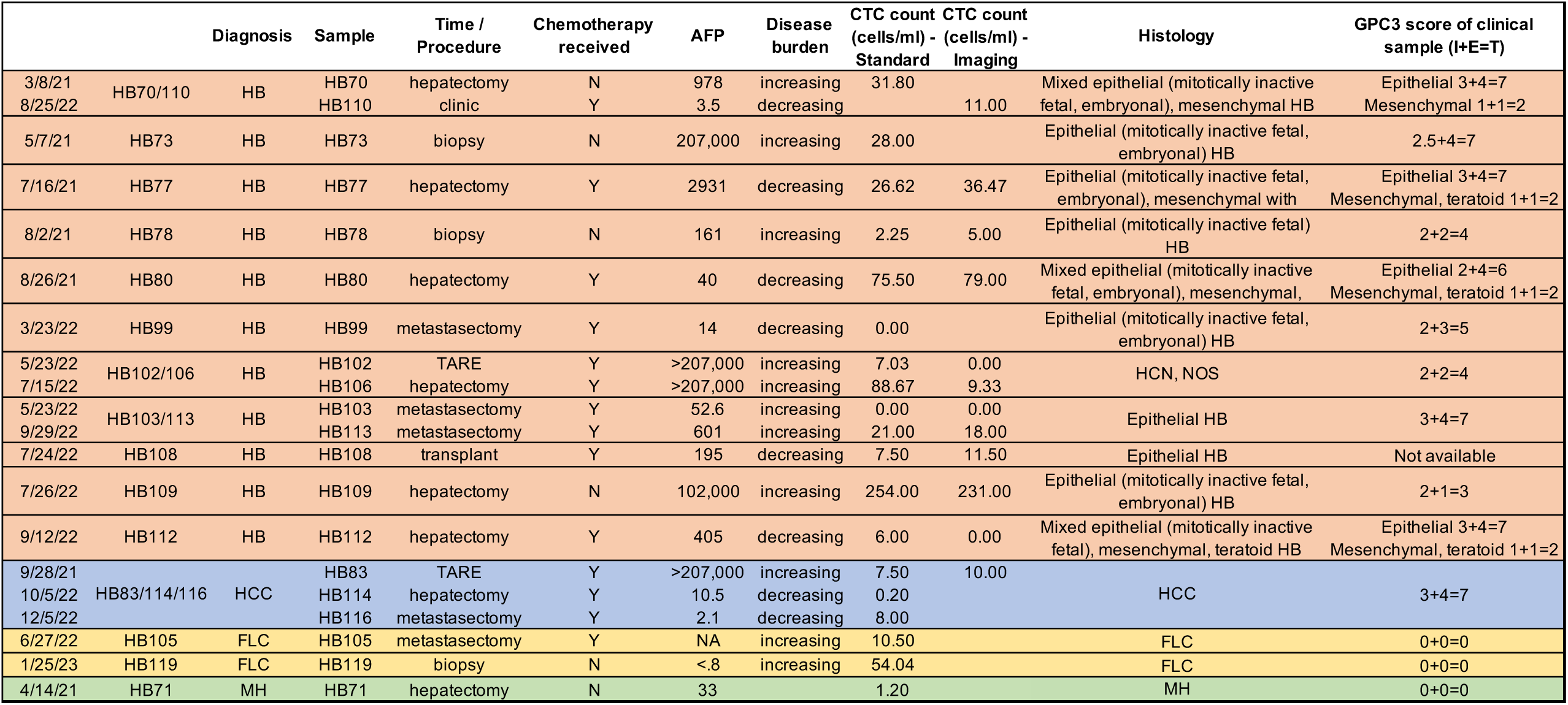
Table of sample data. CTC burden shown is number of DAPI^+^/ICG^+^/GPC3^+^ cells measured by indicated flow cytometry instrument.

For four patients, we had data from at least two patient clinical encounters, and analyses of these samples showed that CTC burden did correlate with overall patient response to therapy. This is shown for one very low risk patient in Figure 4, two high-risk patients in Figure 5, and one HCC patient in Figure 6.

**Figure 4.**
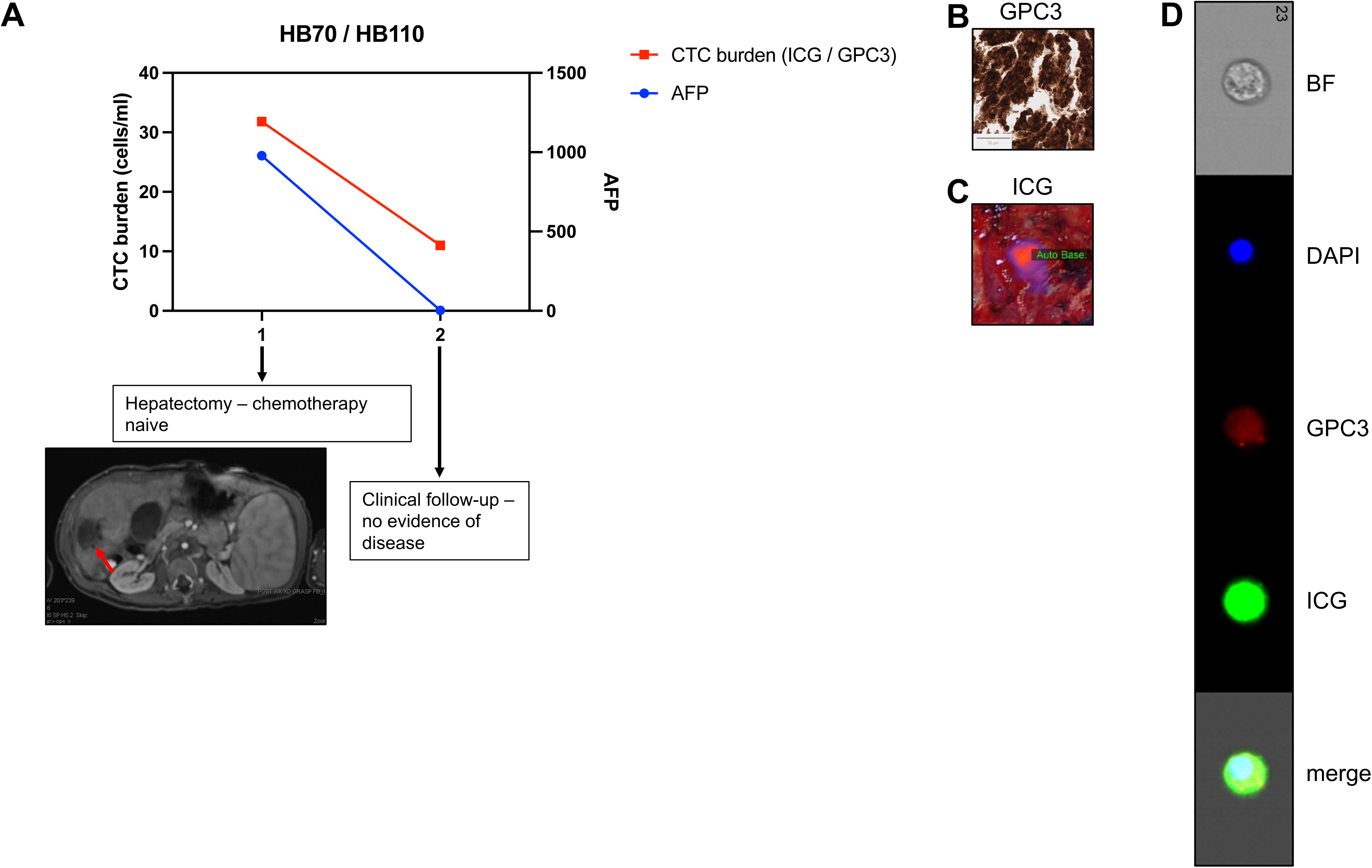
CTC burden in very low risk patient HB70/110. (A) We obtained samples at two time points during the patient’s course of treatment. We analyzed CTC burden after processing whole blood and tagging CTCs with ICG, GPC3, and DAPI, as described. We graphed CTC burden (cells/ml) and serum AFP levels, and both show a drop, correlating with response of the patient to therapy. AFP was assessed by standard clinical tests. CT of primary tumor directly prior to hepatectomy with tumor indicated by red arrow. (B,C) Validation of GPC3^+^ and ICG^+^ primary samples from patient. (B) Histology of primary patient tumor sample showing positivity of sample for GPC3. Scale bar represents 50 μm. (C) Near-infrared imaging of ICG^+^ primary tumor during hepatectomy. (D) Image of ICG^+^/GPC3^+^/DAPI^+^ CTC from Amnis ImageStream instrument.

**Figure 5.**
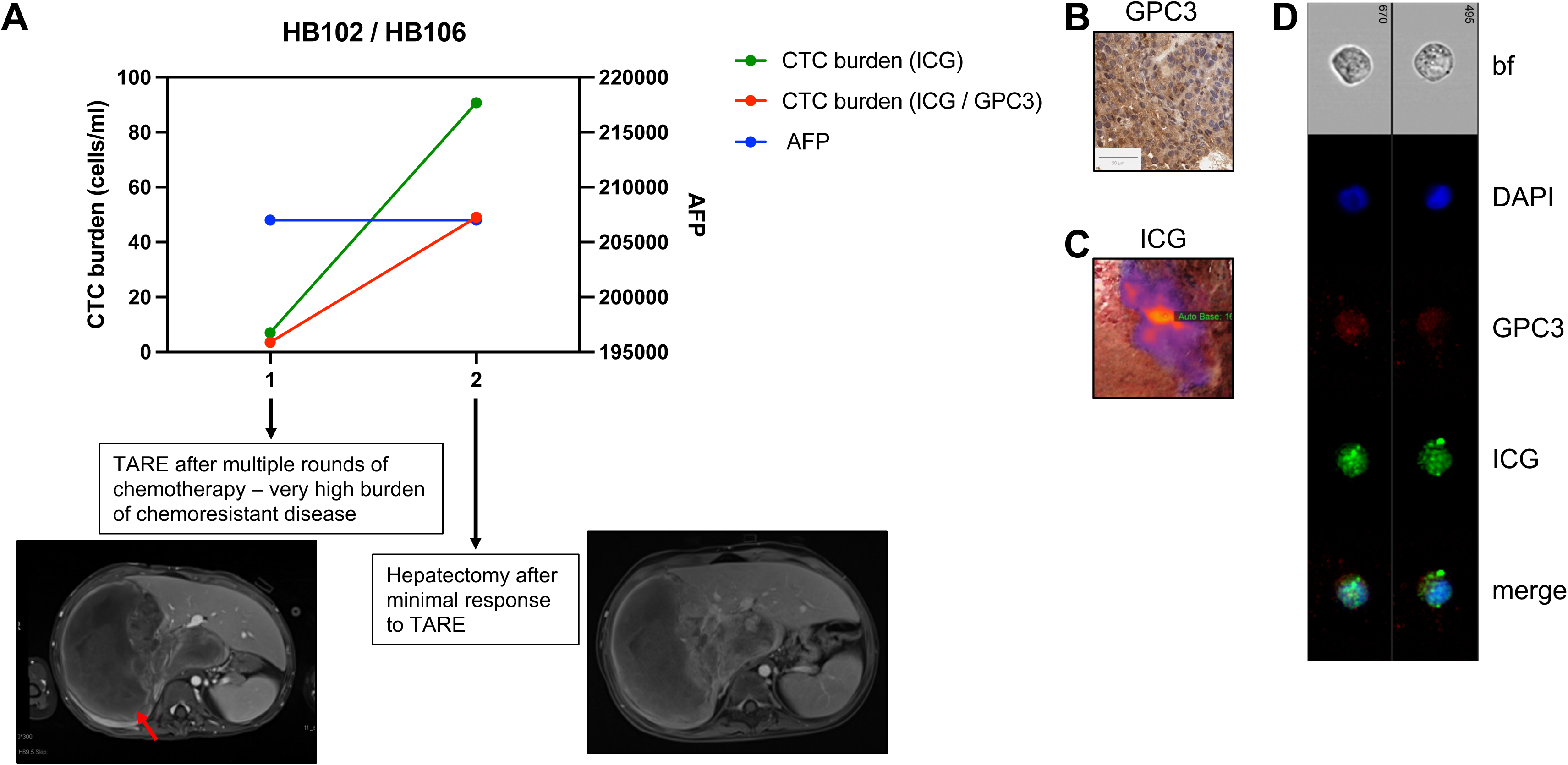
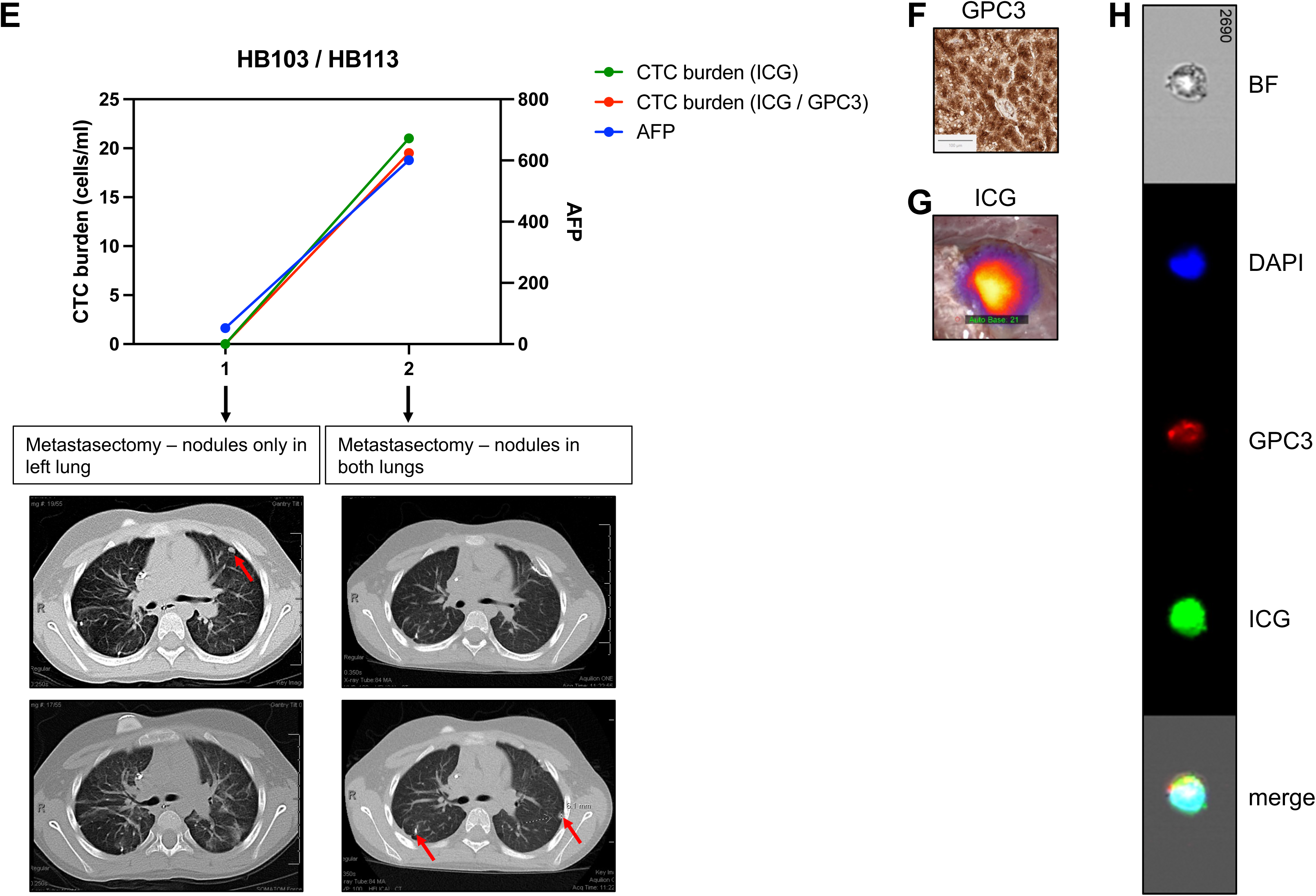
CTC burden in two high-risk patients, HB102/106 (A-D) and HB103/113 (E-H). (A) We obtained samples at two time points during the patient’s course of treatment. We analyzed CTC burden after processing whole blood and tagging CTCs with ICG, GPC3, and DAPI, as described. We graphed CTC burden (cells/ml) and serum AFP levels, and both show an increase, correlating with the patient not responding to therapy. AFP was assessed by standard clinical tests. DAPI^+^/ICG^+^/GPC3^+^ cell number shown is an average of counts by the standard and imaging flow cytometers for both samples. DAPI^+^/ICG^+^ cell number shown was measured by the standard flow cytometer. CT scans from time of TARE and time of hepatectomy showing minimal decrease of tumor (red arrow) size. (B,C) Validation of GPC3^+^ and ICG^+^ primary samples from patient. (B) Histology of primary patient tumor sample showing positivity of sample for GPC3. Scale bar represents 50 μm. (C) Near-infrared imaging of ICG^+^ primary tumor during hepatectomy. (D) Image of two ICG^+^/GPC3^+^/DAPI^+^ CTCs from Amnis ImageStream instrument. (E) We obtained samples at two time points during the patient’s course of treatment. We analyzed CTC burden after processing whole blood and tagging CTCs with ICG, GPC3, and DAPI, as described. We graphed CTC burden (cells/ml) and serum AFP levels, and both show an increase, correlating with the patient not responding to therapy. AFP was assessed by standard clinical tests. DAPI^+^/ICG^+^/GPC3^+^ cell number shown is an average of counts by the standard and imaging flow cytometers for both samples. DAPI^+^/ICG^+^ cell number shown was measured by the standard flow cytometer. CT images shown are same slices in rows from two metastasectomy procedures, showing nodule in left lung at first time point (red arrow, left images) and new nodules present in both lungs (red arrows, right images) at second time point. (F,G) Validation of GPC3^+^ and ICG^+^ primary samples from patient. (F) Histology of primary patient tumor sample showing positivity of sample for GPC3. Scale bar represents 100 μm. (G) Near-infrared imaging of ICG^+^ lung nodule during metastasectomy. (D) Image of ICG^+^/GPC3^+^/DAPI^+^ CTCs from Amnis ImageStream instrument.

**Figure 6.**
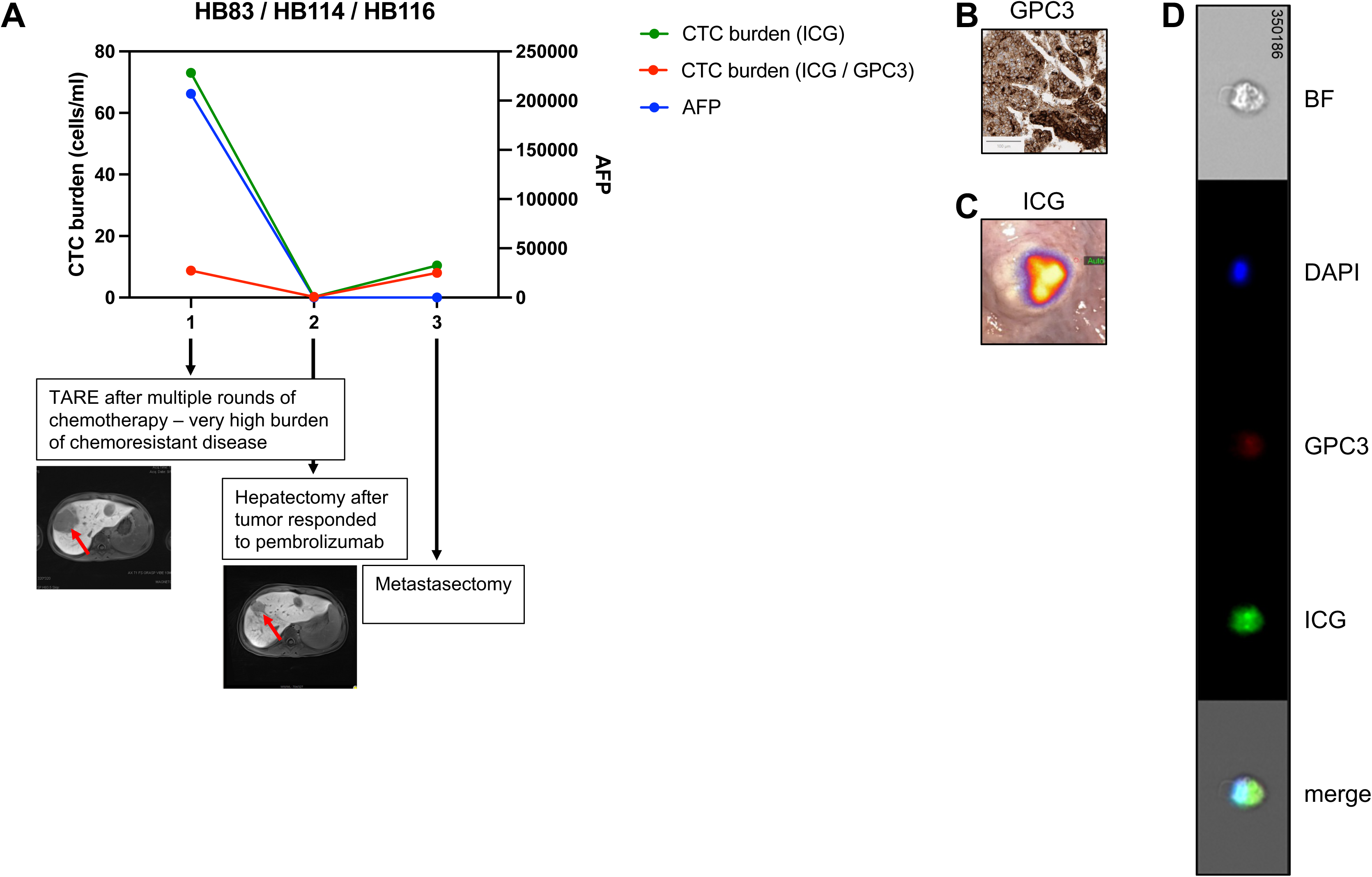
CTC burden in an HCC patient HB83/114/116. (A) We obtained samples at three time points during the patient’s course of treatment. We analyzed CTC burden after processing whole blood and tagging CTCs with ICG, GPC3, and DAPI, as described. We graphed CTC burden (cells/ml) and serum AFP levels, and both show a drop, correlating with response of the patient to therapy. AFP was assessed by standard clinical tests. DAPI^+^/ICG^+^/GPC3^+^ cell number shown is an average of counts by the standard and imaging flow cytometers for the HB83 sample. DAPI^+^/ICG^+^ cell number shown was measured by the standard flow cytometer. CT images from time of TARE and time of hepatectomy with tumor (red arrows) decrease shown. (B,C) Validation of GPC3^+^ and ICG^+^ primary samples from patient. (B) Histology of primary patient tumor sample showing positivity of sample for GPC3. Scale bar represents 100 μm. (C) Near-infrared imaging of ICG^+^ lung nodule during metastasectomy. (D) Image of ICG^+^/GPC3^+^/DAPI^+^ CTC from Amnis ImageStream instrument.

We analyzed two samples from the HB70/HB110 patient, one from time of hepatectomy when the patient had not yet received chemotherapy and one from a follow-up clinical appointment (17.5 months after hepatectomy) when the patient was deemed free of disease (Figure 4). This patient was diagnosed with a very low risk PRETEXT I lesion and was treated with upfront resection followed by two cycles of C5V (cisplatin, 5-fluorouracil, vincristine) adjuvant chemotherapy. Currently, this patient is in remission. This patient was strongly positive for both GPC3 (Table 2, Figure 4B) and ICG (Figure 4C). During that time, CTC burden dropped from 31.8 cells/ml to 11 cells/ml and AFP levels dropped from 978 to 3.5, corresponding to the response of the tumor cells to therapy (Figure 4A). Shown in Figure 4D is an image of a DAPI^+^/ICG^+^/GPC3^+^ CTC analyzed by the Amnis imaging flow cytometer from the sample from the follow-up clinical encounter.

We analyzed two samples from the HB102/HB106 high-risk patient, who was diagnosed with a non-metastatic PRETEXT III HB tumor with HCC features and was treated with three rounds of chemotherapy, followed by transarterial radioembolization (TARE) and then resection. After surgery, the patient received adjuvant pembrolizumab but then died due to disease. We analyzed one sample from when the patient received TARE and one from time of hepatectomy surgery (Figure 5A-D). This patient showed low, heterogeneous GPC3 expression (Table 2, Figure 5B) and strongly positive ICG signal (Figure 5C). During this time, although the patient was receiving chemotherapy, the tumor cells were not responding and the disease burden was increasing. This was assessed based on standard clinical readouts for disease response that were measured during that time. For example, there was no change in AFP level (Figure 5A). In addition, the tumor removed with resection was 30% viable. We also measured an increase in CTC burden from 7.03 cells/ml at time of TARE to 49 cells/ml at time of hepatectomy (Figure 5A). Further, when we reanalyze our flow data for only DAPI^+^/ICG^+^ cells, we see even more CTCs at both time points (Figure 5A). Shown in Figure 5D is an image of two DAPI^+^/ICG^+^/GPC3^+^ CTCs analyzed by the Amnis imaging flow cytometer from the sample from time of hepatectomy.

We also analyzed two samples from the HB103/HB113 high-risk patient who was initially diagnosed with a high-risk PRETEXT IV HB tumor and treated with three cycles of cisplatin/doxorubicin chemotherapy followed by transplant and then completion of carboplatin/doxorubicin chemotherapy. The tumor removed during surgery was >99% viable. The patient was then deemed to be disease free based on no disease shown by imaging and normal AFP levels. After 8 weeks during routine follow-up, the patient was found to have a rising AFP and relapsed metastatic disease by surveillance imaging.

They then underwent six cycles of vincristine/irinotecan and staged resection of lung tumors. During the first metastasectomy surgery, one nodule was identified by CT prior to surgery (Figure 5E, left CT images) and two more were found by ICG during surgery; all three were removed and were positive for disease. During the third metastasectomy surgery, three nodules had been identified by CT (Figure 5E, right CT images) and eight more were identified during surgery by ICG; all 11 nodules were removed and three were positive for tumor. Despite all of this therapy, the patient passed away. We analyzed two blood samples from this patient, both from metastasectomy surgeries after relapse. The patient was also strongly positive for both GPC3 (Table 2, Figure 5F) and ICG (Figure 5G). At the first surgery, the patient only had detectable tumor cells in the left lung (Figure 5E, left CT images) and low AFP of 52.6; at this time point we did not detect CTCs. From the second sample, we detected 19.5 cells/ml, which corresponds to the rise in AFP to 601. We also analyzed the samples for DAPI+/ICG+/GPC3^-^ cells and found that most ICG^+^ cells were also GPC3^+^, consistent with the immunohistochemistry for GPC3 (Figure 5F). Shown in Figure 5H is an image of a DAPI^+^/ICG^+^/GPC3^+^ CTC analyzed by the Amnis imaging flow cytometer from the sample from the second metastasectomy.

We analyzed three samples from the HB83/HB114/HB116 HCC patient who was initially diagnosed with a PRETEXT IV metastatic HB. They received two cycles of cisplatin/5-fluorouracil/vincristine induction chemotherapy and moderately responded with AFP decrease and shrinkage of masses by imaging, including resolution of metastatic lesions in the right lung. They then progressed and underwent a right metastasectomy in efforts to qualify for liver transplant. The metastatic lesion was determined to be HCC, so they were transitioned to HCC-specific therapy and were treated with (1) GEMOX (gemcitabine hydrochloride/oxaliplatin)/sorafenib, (2) TARE, (3) atezolizumab/bevacizumab, (4) IL15-GPC3 CAR T cells, (5) vincristine/irinotecan, (6) lenvatinib, and (7) pembrolizumab. They responded to pembrolizumab, which resulted in a significant AFP decrease and reduction of the primary lesion to allow liver tumor and lung metastasis surgeries. In the metastasectomy, they had a total of 15 lesions removed and only one was still viable for disease. This patient is still alive and receiving treatment. We received three samples from this patient, the first from time of TARE after multiple rounds of chemotherapy when there was high disease burden, the second from hepatectomy after the patient responded to phase 1 experimental therapy, and the third at time of metastasectomy (Figure 6A). This patient was strongly positive for both GPC3 (Table 2, Figure 6B) and ICG (Figure 6C). The first sample showed higher CTC burden of 8.75 cells/ml, corresponding to the very high AFP of >207,000 (Figure 6A). The tumor responded to therapy as indicated by standard clinical measures, and the CTC burden dropped to 0.2 cells/ml and AFP to 10.5 (Figure 6A). At the third time point, the CTC count rose to 8 cells/ml while the AFP continued to drop to 2.1 (Figure 6A). We also analyzed the samples for DAPI+/ICG+/GPC3^-^ cells and found that there were many ICG^+^ cells that were GPC3^-^ in the first sample, which is not consistent with the GPC3 immunohistochemistry (Figure 6A). Shown in Figure 6D is an image of a DAPI^+^/ICG^+^/GPC3^+^ CTC analyzed by the Amnis imaging flow cytometer from the first sample during TARE.

## DISCUSSION

The major challenge in the field of CTC biology has been how to unambiguously identify these rare cells for research and clinical purposes. Initial work used epithelial markers to distinguish CTCs that arose from carcinomas. In fact, the only FDA-approved method for CTC enumeration for clinical care is the CELLSEARCH platform, which relies on such tumor markers to identify CTCs^31^. However, it has become clear more recently that CTCs from carcinomas may undergo epithelial mesenchymal transition as they intravasate into vessels, thus losing their epithelial markers^32,33^. Thus, other unbiased methods have emerged, including isolation of CTCs based on physical properties such as size, charge, density, and elasticity^33^. Our work using a fluorescent dye that is specifically accumulated in liver tumor cells addresses this challenge. This use of a dye that stains all tumor cells is particularly important to pediatric liver cancer, which presents as a very heterogenous tumor.

The second overarching problem to address is how to integrate CTC numbers into clinical care. Most work on evaluation of outcomes associated with CTCs has been focused on common adult cancers including colorectal, breast, and genitourinary tumors^18,34–36^. A key clinical trial with 778 patients with hormone receptor-positive, ERBB2-negative metastatic breast cancer (STIC CTC METABREAST trial) showed that separating patients into chemotherapy or endocrine therapy based on CTC count instead of physician’s choice improved median progression free survival by 5.6 months^37^. By and large, these studies have established that the presence of CTCs is associated with worse prognosis in adult patients and that patients with elevated CTC count will likely benefit from more aggressive therapies. With these more common adult solid tumors, large numbers of patients can be more quickly evaluated to draw clear conclusions about how numbers of CTCs can predict therapy responses and outcomes. There is not yet enough data focused on rare tumors, such as pediatric solid tumors, to inform use of CTC numbers clinically with these patients. In the area of pediatric solid tumors, major liquid biopsy work has focused on neuroblastoma and sarcoma^38^. While both cancers have established worse outcomes with the presence of CTCs, most studies have instead focused on outcomes associated with ctDNA^38^. Given that the presence of CTCs has been well established to be associated with poor prognosis in many other solid tumor types, our liquid biopsy protocol may provide this bridge to clinical translation for pediatric liver tumors. In our small study, we are not able to correlate CTC numbers with any patient characteristics or outcomes, likely because our patients represent a range of diagnoses, stages and risk groups, and other disease attributes. Continuation of this study with additional patients will allow us to examine CTC counts in all patients at, for example, diagnosis or relapse to see if that number predicts outcomes for pediatric liver cancer.

Our data most convincingly shows that changes in CTC burden overtime while a patient is receiving therapy correlates with the patient’s response to therapy and may even give additional information that is not available from other clinical assays. Notably, CTC burden seems to be a better readout of response for the HCC patient (HB83/114/116, Figure 6) because this patient showed an increase in CTCs between hepatectomy and metastasectomy procedures when it was clear they still had disease present by imaging; during this same timeframe, AFP levels for this patient dropped from 10.5 to 2.1. For patient HB102/106, we saw a rising CTC burden between TARE and hepatectomy procedures when tumor burden was also rising; during this same timeframe, AFP levels for this patient had already reached the maximum measurable level and, therefore, provided no further information about response. Taken together, this data shows that the best utilization of our test is to compare CTC burdens in sequential blood draws from the same patient during therapy to show whether tumors are responding. Continuation of our work will focus on collecting and analyzing diagnostic blood samples for baseline information about patients that we can compare further counts with during the course of therapy.

Importantly, analyses of ICG and GPC3 positivity from intraoperative imaging and histology, respectively, validated and informed our use of ICG and GPC3 to identify CTCs. All patients tested were strongly positive for ICG, confirming our use of ICG as the major identifier of liver tumor identity. By and large, patients that showed low or heterogenous GPC3 expression also showed the presence of CTCs that were DAPI^+^/ICG^+^ but GPC3^-^. This data shows that the crosstalk between surgeons and pathologists is key for informing how our CTC panel is analyzed. Future work can focus on substituting additional markers for GPC3 for GPC3^-^ tumors, particularly markers that may be more specific to FLC. This work also shows that our test can still be consistently used even in the absence of GPC3 expression. In our initial experiments to examine ICG accumulation in liver tumor cells, we also see residual ICG signal in the lung cancer cell line A549 by flow cytometry, which is interesting given evidence that lung cancers accumulate ICG^39,40^. Our use of ICG to identify CTCs may be more widely applicable to diverse tumors that accumulate ICG, and our panel could be easily adapted to these tumors with substitution of an alternative tumor-specific validation marker instead of GPC3.

In our validation of our panel with non-cancer control samples, as well as one blood sample from a non-malignant MH tumor, we show that this assay detects no more than 1.2 cells per ml of blood, proving the specificity and sensitivity of the test. This presence or absence of CTCs is invaluable information that can inform whether a patient is at risk for metastasis. However, this is complicated by the established fact that CTCs have a short half-life^41^ and not all CTCs are capable of giving rise to detectable metastatic disease^33^. Future work in our lab and others is focused on how to identify and target these “bad” CTCs that are capable of colonizing organs outside of the organ of origin, and this information will also inform markers for more specific CTC tests to identify the most aggressive CTCs.

Taken together, this work shows a novel way to identify CTCs with the far-red fluorescent dye ICG that is already used clinically and is a compelling example of bedside-to-bench scientific research. Notably, our test can be used for the two most common malignant pediatric primary tumors of the liver, HB and HCC, as both tumor cell types accumulate ICG. This work will form a foundation for future work to establish how CTC number can be used in the routine clinical care of these patients.

## Abbreviations

HB: hepatoblastoma
HCC: hepatocellular carcinoma
CTC: circulating tumor cell
ICG: indocyanine green
GPC3: Glypican-3
OS: overall survival
FLC: fibrolamellar carcinoma
AFP: Alpha-fetoprotein
ctDNA: circulating tumor DNA
CAR: chimeric antigen receptor
ATCC: American Type Culture Collection
EMEM: Eagle’s Minimum Essential Medium
FBS: fetal bovine serum
PDX: patient-derived xenograft
STR: short tandem repeat
FSC: forward scatter
SSC: side scatter
IRB: Institutional Review Board
PRETEXT: pretreatment extent of disease
COG: Children’s Oncology Group
VI: vascular invasion
MH: mesenchymal hamartoma
TARE: transarterial radioembolization

## Acknowledgements

This project was supported by the Cytometry and Cell Sorting Core at Baylor College of Medicine with funding from the CPRIT Core Facility Support Award (CPRIT-RP180672), the NIH (CA125123 and RR024574) and the assistance of Joel M. Sederstrom.

