## Supplementary figures and images for "An indocyanine green-based liquid biopsy test for circulating tumor cells for pediatric liver cancer"

### Supplementary Figure 1

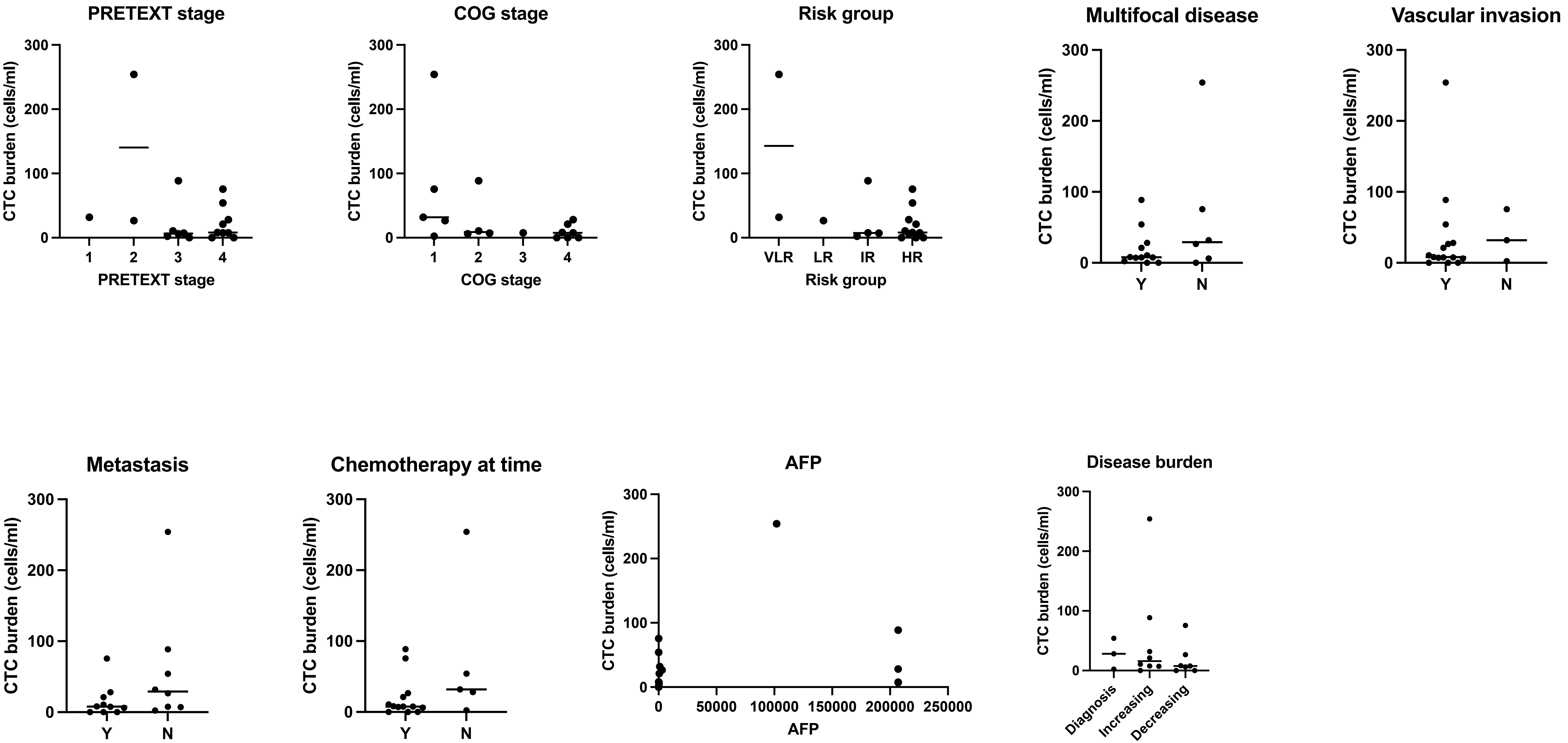


Supplementary Figure 1
